# Multifactorial Dynamics of White Matter Connectivity During Adolescence

**DOI:** 10.1101/215152

**Authors:** Birkan Tunç, Drew Parker, Russell T. Shinohara, Mark A. Elliott, Kosha Ruparel, Raquel E. Gur, Ruben C. Gur, Ragini Verma

**Affiliations:** Center for Biomedical Image Computing and Analytics, Department of Radiology; Department of Biostatistics and Epidemiology; Center for Magnetic Resonance and Optical Imaging, Department of Radiology; Neuropsychiatry Section, Department of Psychiatry Perelman School of Medicine, University of Pennsylvania, Philadelphia, PA 19104, USA.

## Abstract

Studying developmental changes in white matter connectivity is critical for understanding neurobiological substrates of cognition, learning, and neuropsychiatric disorders. This becomes especially important during adolescence when a rapid expansion of the behavioral repertoire occurs. Several factors such as brain geometry, genetic expression profiles, and higher level architectural specifications such as the presence of segregated modules have been associated with the observed organization of white matter connections. However, we lack understanding of the extent to which such factors jointly describe the brain network organization, nor have insights into how their contribution changes developmentally. We constructed a multifactorial model of white matter connectivity using Bayesian network analysis and tested it with diffusion imaging data from a large community sample. We investigated contributions of multiple factors in explaining observed connectivity, including architectural specifications, which promote a modular yet integrative organization, and brain’s geometric and genetic features. Our results demonstrated that the initially dominant geometric and genetic factors become less influential with age, whereas the effect of architectural specifications increases. The identified structural modules are associated with well-known functional systems, and the level of association increases with age. This integrative analysis provides a computational characterization of the normative evolution of structural connectivity during adolescence.

## Introduction

The human brain, as depicted by a network of interconnected regions, acquires a distinctive structural and functional architecture over the course of development [1], [2]. This distinctive architecture has a hierarchical modular organization [3], with special hub regions that are theorized to facilitate the integration of modules [4], [5]. It is generally postulated that the structural organization of the brain provides a basis for the emergent functional systems [6] and thereby, of cognitive capacities [7]. Therefore, studying developmental changes in the white matter connectivity and resulting structural organization is critical for understanding neurobiological substrates of cognition [8], [9], learning [10], and psychiatric conditions [11]. This becomes especially important during adolescence and young adulthood when the human brain undergoes a protracted course of remodeling to support the rapid expansion of its behavioral repertoire [12].

Various empirical studies have revealed several factors that are correlated with the observed white matter connectivity, including regional genetic expression profiles, brain geometry, and wiring costs of fibers [13]–[16]. Despite the significant correlation of these factors with the observed structural connectivity, they do not fully explain the distinctive organization of the brain network [14], [17]. For this reason, in addition to such intrinsic factors, several higher level factors, corresponding to certain architectural specifications (e.g. presence of segregated modules), have been also considered to explain distinctive features of the brain network organization [18]. All these findings highlight the need for a robust multifactorial analysis of the white matter connectivity. This multifactorial analysis is crucial to understand the extent to which multiple factors, such as brain geometry and architectural specifications, jointly describe the brain network organization. Such a multifactorial analysis can also reveal the temporal changes in the contribution of these factors during development, a topic that has been only sparsely studied.

In this work, we have developed a multifactorial generative model of structural connectivity using Bayesian network analysis [19], [20]. This approach facilitates elucidating the contribution of different factors in explaining the empirical connectivity in a data-driven fashion, without necessarily assuming any causal theories regarding the formation of the network. One important advantage of generative models is that developmental effects can be studied by analyzing the evolution of model parameters across ages.

Using our generative model and diffusion imaging data of a large community sample of youth, collected as a part of The Philadelphia Neurodevelopmental Cohort (PNC) dataset [11], we studied the contribution of certain architectural specifications, which promote a modular yet integrative architecture, and brain’s geometric and genetic features in explaining the observed structural connectivity. First, we investigated how much of the observed structural connectivity can be described by a base model of connectivity that includes only geometric and genetic factors. Then, we demonstrated how the inclusion of the architectural specifications increases the accuracy of the generative model in explaining the connectivity. Finally, we quantified the developmental effects in our multifactorial model in terms of the relative contributions of the factors across ages.

Our results demonstrated that the initially dominant geometric and genetic factors become less influential with age, whereas the effect of higher level architectural specifications that promote a modular yet integrative organization increases. The identified structural modules of the brain are associated with well-known functional systems, and the level of association increases with age as well. This integrative analysis provides a computational characterization of the normative changes in structural connectivity in the human brain during the course of development.

## MATERIALS AND METHODS

### Participants

Institutional Review Board approval was obtained from the University of Pennsylvania and the Children’s Hospital of Philadelphia. We used a large sample of healthy young individuals (ages between 8-22 years) from the Philadelphia Neurodevelopmental Cohort (PNC) dataset [11], each assessed using diffusion MRI. Participants were excluded due to poor imaging data quality, or a history that suggested potential abnormalities of brain development such as medical problems that might affect brain function, inpatient psychiatric hospitalization, or current use of psychotropic medication. The final study sample included 818 participants, with 361 males (mean age: 15.10 years, std: 3.44) and 457 females (mean age: 15.25 years, std: 3.33).

### Image Acquisition and Brain Network Construction

Diffusion weighted magnetic resonance imaging (dMRI) scans were acquired for each individual. Quality assurance of the data was conducted to detect motion and scanner related artifacts and outliers, using the procedure described in [21]. The details of image acquisition and network construction are given in **Supplementary Note S1** and illustrated in **Fig. 1**. Our pipeline included brain extraction using FSL [22], DWI de-noising using Slicer [23], tensor fitting using multivariate linear fitting [24], cortical/subcortical segmentation using Freesurfer [25], and probabilistic tractography (seeded from white matter / gray matter boundary) using the probtrackx utility of FSL [22].

**Figure 1.**
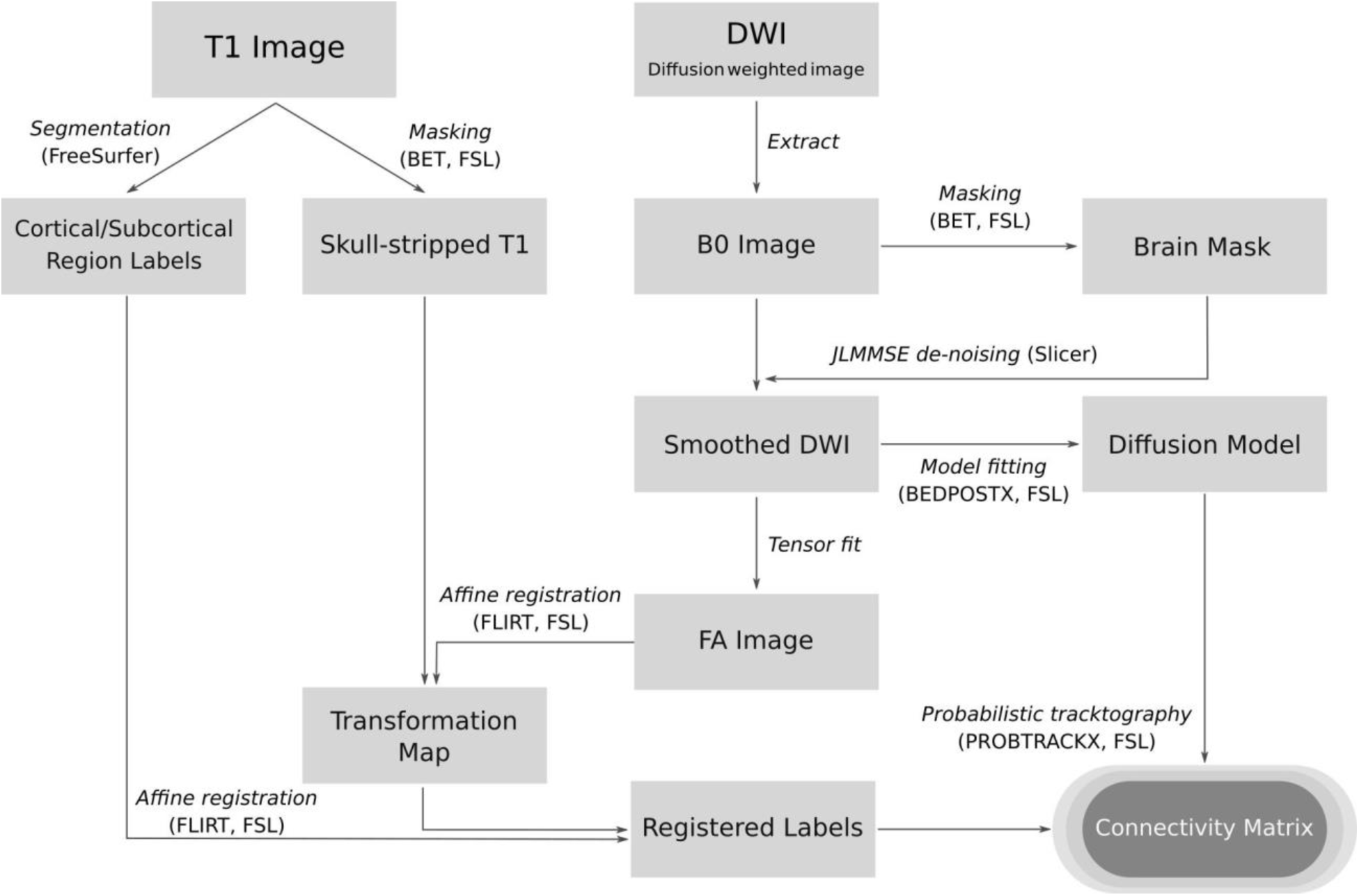
Image processing pipeline. The steps of the image processing and connectivity matrix construction pipeline are displayed as arrows between the rectangle boxes that represent inputs and outputs of the steps. The software tools used are given in parentheses.

The final brain network of a participant had 86 nodes corresponding to the gray matter regions of interest (ROIs), including 34 cortical regions of the Desikan atlas [26], 8 subcortical regions, and cerebellum in the left and right hemispheres [25]. The complete list of regions is given in **Supplemental File 1**. The edges of the network had weights corresponding to the normalized number of streamlines between regions, as generated using probabilistic tractography. The edge weight between two regions i and j was calculated as o_ij_ = (w_ij_ + w_ji_)/2, where w_ij_ = s_ij_/v_i_, s_ij_ is the streamline count between the regions, and v_i_ is the volume (the number of voxels used for seeding) of the region i. The normalization by the region volume accounts for the differences in the region size. The final value was rounded to have integer valued edge weights.

In order to prune possibly spurious connections, we used consistency based thresholding [27], keeping only the most consistent edges across all participants. Due to broadly ranging estimates of actual connection density in the literature [5], [27], [28], we repeated our experiments in a density range of 10-30%, with different thresholds. Here, we report results for 15% density. Very consistent results were achieved for other density levels (results are provided in **Supplementary Notes S8**).

### Generative Model for Brain Connectivity

A schematic illustration of the generative model is shown in **Fig. 2**. Given a set of prior expectations (**E**) over the streamline counts between N regions, we model the observed streamline counts (**O**) between the regions by a Dirichlet-Multinomial distribution, as proposed before [29], [30].

**Figure 2.**
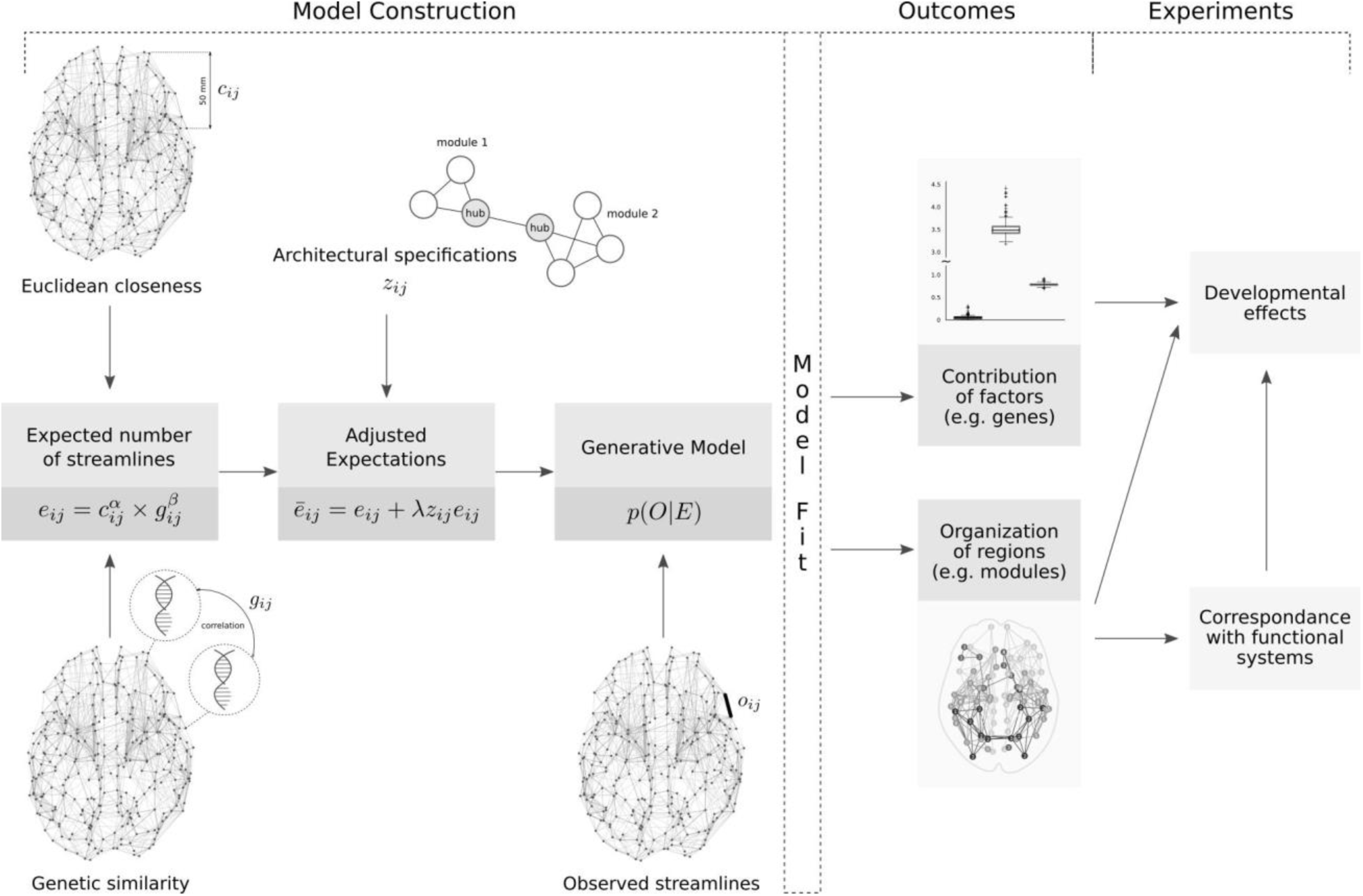
Illustration of methodology and experiments. The generative model of streamline connectivity is constructed, using the expected number of connections and the observed streamline counts as the input. The expected number of connections are calculated based on the genetic and geometric (Euclidean closeness) relationships among regions, as well as by considering certain architectural specifications. After model fitting, the two main outcomes are the quantification of the contribution of different factors in explaining the observed streamline counts and the organization of brain regions. Finally, a developmental analysis to probe changes in these outcomes across ages is performed.

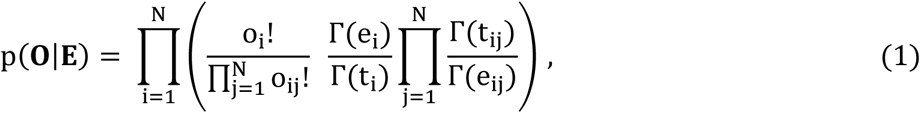

where 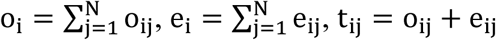, and 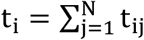. The variables o_ij_ and e_ij_ represent the observed number of streamlines (o_ij_) between regions i and j, and our prior expectations (e_ij_) for the same.

In this study, we propose a novel base model of streamline connectivity by postulating specific expectations (e_ij_) for the number of streamlines between regions, relying on their geometric and genetic features. We considered different mathematical expressions for modeling e_ij_ and selected the following one using Bayesian model comparison [31] (see **Supplementary Note S2** for details).

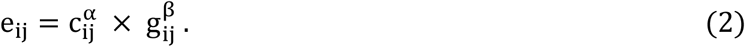

In the base model, c_ij_ corresponds to the Euclidean closeness (inverse of distance) of region centers, and the second factor g_ij_ is the genetic similarity between regions. The unknown exponents, α and β, determine the contribution of each factor.

In our experiments, the first factor c_ij_ was different for each participant, calculated from their T1 images. The second factor g_ij_, however, was fixed for all participants, calculated using Allen Institute for Brain Science (AIBS) microarray expression dataset [32], as explained below. In order to facilitate a comparison between the contribution of different factors, both c_ij_ and g_ij_ were normalized to have the same maximum value with the observed number of streamlines (o_ij_), for each participant individually. Then, the resulting matrix **E** ≡ {e_ij_} was normalized so that its summation is equal to the summation of observations **O** ≡ {o_ij_}. This guarantees that the maximum likelihood solution of **Eq. 1** is attained when e_ij_ = o_ij_.

Although the Dirichlet-Multinomial distribution is a natural choice for modeling relative strength of connectivity among regions and has been commonly used in the literature for similar purposes [20], [29], [30], the plausibility of our parametric model needs to be established prior to subsequent experiments. For this reason, we demonstrated that our generative model performs very similar to nonparametric models that have been developed for specialized purposes (see **Supplementary Note S3** for details).

### Genome-wide Expression Profiles

AIBS microarray expression dataset [32] consists of 3702 brain tissue samples from brains of six donors (5 males, 1 female, ages:24-57 years) [33]. Each sample is assessed by 58,692 probes. The expression levels were normalized across brains during comprehensive pre-processing steps [34], [35]. Among all probes available in the microarrays, we only considered the 17,348 uniquely annotated transcripts that are included in [36]. In order to select probes corresponding to the genes that are most relevant to brain function, we used the highest 10% of genes ranked by their differential stability (DS) indices [36].

Please refer to **Supplementary Note S4** for details on our pipeline to calculate the genetic similarity between regions (g_ij_). Our pipeline included steps to find correspondence between AIBS tissue samples and 86 regions of our atlas, to exclude genes with expression values that were not significantly different than the background expression level, to decrease the dimensionality using linear discriminant analysis (LDA), to calculate Pearson correlation between regional genome-wide expression profiles, and to adjust the genetic similarity values for distance between regions.

### Detection of Modules and Hubs

Architectural specifications, corresponding to the presence of a modular structure and integrative hub regions, were imposed by modifying e_ij_ as

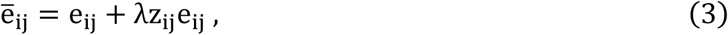

where the structure of the variable z_ij_ encodes the architectural specifications. When assuming only modules but not hubs, z_ij_ is 1 if two regions are in the same module and −1 otherwise, and e_ij_ is calculated as in **Eq. 2**. Each module defines a cluster of regions that are densely connected to each other while only being sparsely connected to the regions outside their modules [37]. This model increases the number of expected streamlines between any two regions by λe_ij_, if regions are in the same module, while decreasing the same if they are in different modules. When there are both modules and hubs, z_ij_ is 1 if two regions are in the same module, or at least one of them is a hub region and they are in the same hemisphere, and −1 otherwise. In this case, the model expects hub regions to make more connections with all other regions. The same hemisphere constraint was used to relax this high demanding expectation.

### Inference of Model Parameters

The unknown parameters α, β, and λ in **Eq. 2**,**3** and the variable z_ij_ in **Eq. 3** were estimated by maximizing the model likelihood in **Eq. 1**. This was performed for each participant individually. We estimated the unknown parameters for the three scenarios (base model, modular model, and modular yet integrated model) independently, in order to compare between the scenarios. Please refer to **Supplementary Note S5** for implementation details.

### Proportion of Explained Connectivity

For each participant, we calculated three likelihood values, L, L_r_, L_e_, corresponding to the likelihood of the model with expectations defined (1) by **Eq. 2** or **Eq. 3**, (2) by the random model with e_ij_ = constant, and (3) by the empirical model with e_ij_ = o_ij_ (observed streamline counts), respectively.

The random model (L_r_) defines the expectations based on pure chance, whereas the empirical model (L_e_) is the maximum value that our generative model can reach. Thus, the measure

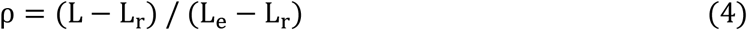

can be considered as a measure of the proportion of the observed connectivity that is explained by the generative model. Note that this measure has an upper bound of 1, and is comparable across participants. The contribution of different factors (e.g., distance or genetic similarity), can thus be quantified by calculating ρ values for models including single factors, or by comparing ρ values with and without using a specific factor in the model.

### Statistical Significance of Factors

In order to calculate the statistical significance of the contribution of different factors (e.g., closeness or genetic similarity), we used the Wilcoxon signed-rank test to compare the Akaike Information Criterion (AIC) [31] values of two models with and without the selected factor. The null hypothesis assumes that difference between the two models follows a symmetric distribution around zero. Additionally, in order to assess the significance of identified modules, compared to randomly defined ones, we used permutation testing. The null hypothesis assumes that an increase in ρ value (**Eq. 4**), at least as high as the actual one, can be achieved by random assignment of regions into similarly sized modules. Details are provided in **Supplementary Note S6**.

### Consensus Modules

The generative model of streamline counts (**Eq. 1**) was run individually for each participant. Therefore, the modular organization (that is, the number of modules and assignment of regions into modules) shows differences between participants. In order to define a consensus of modular organization, we used the mean brain network (o_ij_) and the mean closeness map (c_ij_) of all participants, and estimated the unknown parameters α, β, λ, **m**_i_, **h** again by maximizing the model likelihood in **Eq. 1**. Note that genetic factor g_ij_ was already fixed for all participants.

### Functional Systems

In order to study correspondence between identified structural modules and functional systems of the brain network, we assigned the 86 brain regions to 10 functional systems [38], namely auditory, cingulo-opercular, default mode, dorsal attention, frontoparietal, motor, subcortical, ventral attention, visual, and others, using the definitions from Gu et al. 2014 [39] (see **Supplementary File 1** for the complete list). This resulted in predefined functional systems that are common for all participants.

In order to quantify the similarity between the alliance of regions determined by consensus structural modules and the functional system, we used Normalized Mutual Information (NMI) [40]. NMI quantifies the agreement between structural modules and functional systems in terms of the assignment of regions to same/different modules/systems. It has values between 0 and 1, where 1 indicates perfect agreement. Statistical significance of correspondence between structural modules and functional systems was assessed using permutation testing. Please refer to **Supplementary Note S7** for details.

## RESULTS

### Plausibility of the Generative Model

We compared the networks simulated by our generative model with those that were generated by a non-parametric model [14]. Our model was able to better represent the observed streamlines counts as compared to the nonparametric model, while both models generated networks with similar topological features. Results are shown in **Supplementary Fig. S2**. We also compared the modules identified by our generative model to those that were identified by Louvain method for community detection [41]. We first detected modules using Louvain method and then detected the same number of modules using our model. There was a substantial agreement between identified modules. Results are shown in **Supplementary Fig. S3**.

### Base Model of Streamline Connections

We first defined a base model of connectivity that includes only geometric and genetic factors. Illustration of the two factors (**Fig. 3a**), namely physical closeness and genetic similarity (c_ij_, g_ij_) provides an initial qualitative assessment of the contribution of the factors in explaining observed streamline counts between regions. According to **Fig. 3a**, physical closeness seems to be the dominant factor, with the overall organization of the observed connectivity matrix being very similar to the organization of the adjacency matrix defined by physical closeness. The most important specification that the genetic factor provides is the distinction between cortical, subcortical, and cerebellar regions. A closer look into the genetic similarity, however, reveals that the genetic similarity between regions is not homogenous inside the cortex, reflecting distinctions between different lobes.

**Figure 3.**
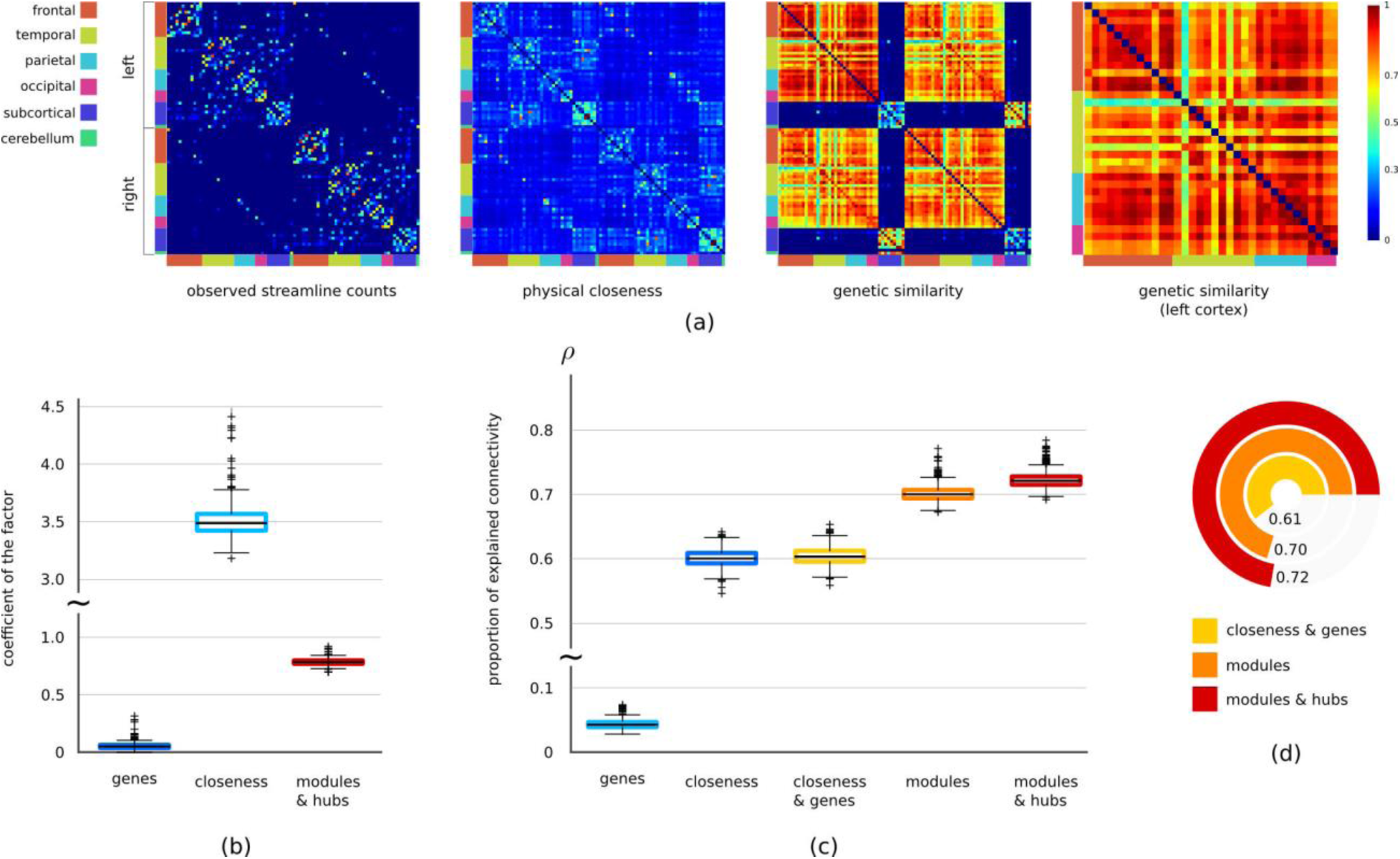
Contributions of multiple factors in explaining observed structural connectivity. (a) Illustration of connectivity matrices encoding observed streamline counts, physical closeness of regions, and genetic similarity between region. The matrices depict the strength of connectivity between regions, as measured by different factors. Rows and columns correspond to regions in the same order. Connectivity matrices are averaged across all participants and normalized to the range 0-1 for visualization purposes. (b) Coefficients (α, β, λ) of different factors in the generative model. Box plots summarize values across all participants. (c) The proportion of explained connectivity (ρ) by models including different sets of factors. The models including modules and hubs also include closeness and genetic similarity factors. All pairwise comparisons between models using Wilcoxon signed-rank test yield statistically significant differences. (d) Comparisons of mean ρ values between three models, namely the base model (closeness and genes), the modular model, and the modular yet integrative model.

Note that for all following analyses, the generative model was run individually for each participant, and summary statistics were computed. The base model was run to infer participant-specific parameters (α, β, see **Eq. 2**) and to compute model likelihoods. We then computed the proportion of the observed connectivity that is explained by the model (ρ, see **Eq. 4**). The measure ρ quantifies the improvement over a random model and has a maximum value of 1.

Our experiments revealed that the most dominant factor in describing streamline counts between regions was physical closeness (**Fig. 3b, c**). When using single factors alone, we had average (across all participants) ρ of 0.60 and 0.05 for closeness and genetic similarity, respectively. Inclusion of the closeness factor increased ρ by 0.56 on average (95% CI: 0.559, 0.561) as compared to the model including only the genetic factors (Wilcoxon signed-rank test to compare AIC values, p < 1 × 10^−16^). The average increase in ρ was 0.003 (95% CI: 0.002, 0.004) with the inclusion of the genetic factor in the model including only closeness factor (Wilcoxon signed-rank test to compare AIC values, p < 1 × 10^−16^). Finally, when using the both factors (**Fig. 3c, d**), we observed the best model fit (ρ=0.61). Note that inclusion of a factor does not increase ρ in a linear fashion due to nonlinear dependencies between factors and the resulting model likelihood.

### Architectural Specifications of Brain Network

With the inclusion of the architectural specifications that encodes the modular structure, we observed a significantly better model fit (ρ=0.70) (**Fig. 3c, d**), with average 0.096 increase in ρ (95% CI: 0.096, 0.098) compared to the base model (Wilcoxon signed-rank test to compare AIC values, p < 1 × 10^−16^). We also assess the significance of modules compared to randomly defined modules (permutation testing with random modules, p < 1 × 10^−3^).

After imposing both modular structure and presence of hub regions, the average ρ was 0.72, which was a significant increase compared to the modular structure alone and to the base model (Wilcoxon signed-rank test to compare AIC values, p < 1 × 10^−16^) (**Fig. 3c**). Thus, our analyses supported the hypothesis that architectural specifications have a significant role in explaining the observed streamline counts between regions.

### A Modular yet Integrative Organization

We identified 6 consensus structural modules. Two pairs of symmetric modules and two modules extending to both hemispheres in **Fig. 4a, b** reflect the consensus of module assignments from all participants. The final assignment of regions into modules is given in **Supplemental File 4**. The consensus set of hub regions for all participants is illustrated in **Fig. 4c**.

**Figure 4.**
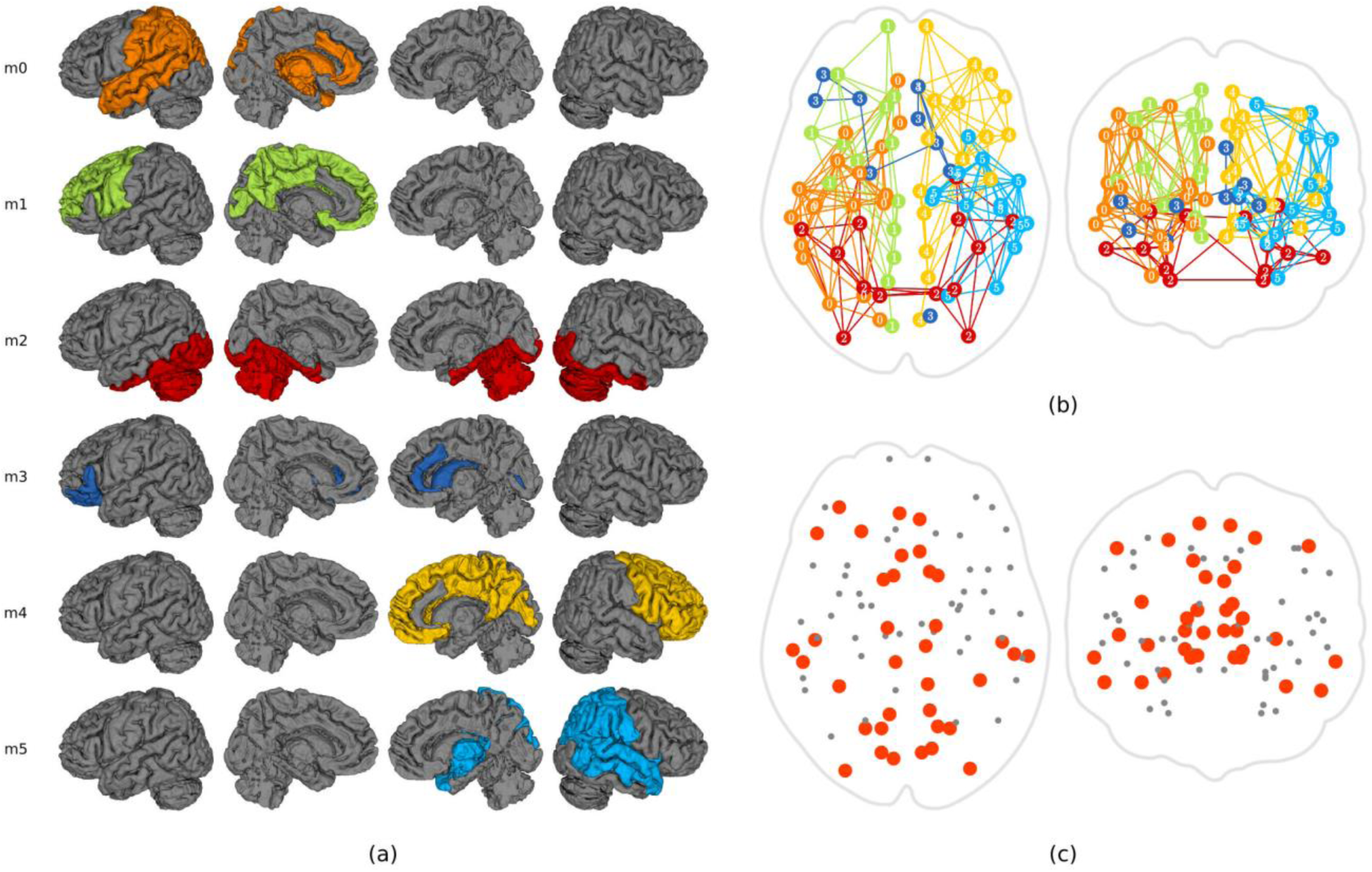
Modules and hubs of the structural brain network. (a) 6 modules were identified with two bi-hemispheric (modules 2,3) and two symmetric pairs of modules. The alliance of regions into modules reflects the functional correlates of these modules. For instance, module 2 mainly consists of regions of the visual system. The modules 1 and 4 form a structural basis for several functional systems including the default mode network, cingulo-opercular system, and ventral attention system. Connections between regions of the module 0 constitute an integral part of the language processing system. (b) Visualization of same modules on axial and coronal slices. (c) Hub regions of the brain network. Hubs are mainly accumulated at the anterior, posterior, and temporal regions instead of being around the geometric center of the network that would be preferred by a geometric model of connectivity.

Additionally, we investigated how structural network organization is related to the functional specialization of regions. We calculated the similarity between the alliance of regions determined by the structural modules and the predefined functional systems using Normalized Mutual Information (NMI). This yielded an NMI value of 0.48 (permutation testing with random modules, p = 0.0003), suggesting a significant functional correlate of structural modules.

### Developmental Effects

Participant-specific parameters (α, β, λ) and ρ values were studied across ages in order to show how model parameters and the contribution of different factors evolve during development. When the full model was run with the architectural specifications, we observed significant changes in the model parameters with age. The coefficient of the physical closeness (α) showed a significant developmental decrease (Pearson correlation coefficient r = −0.18, p = 5 × 10^−7^), while the coefficient of the genetic similarity (β) showed a significant increase (Pearson correlation coefficient r = 0.12, p = 0.0003). We also observed a significant increase in the parameter (λ) encoding architectural specifications corresponding to a modular yet integrative organization (Pearson correlation coefficient r = 0.26, p = 5 × 10^−14^).

We observed a decreasing joint contribution of physical closeness and genetic similarity in the observed streamline counts (**Fig. 5a, b, c**). The proportion of explained connectivity (ρ) showed a significant decrease by age when using only genetic similarity (Pearson correlation coefficient r = −0.14, p = 5 × 10^−5^), only physical closeness (Pearson correlation coefficient r = −0.26, p = 4 × 10^−14^), and both (Pearson correlation coefficient r = −0.26, p = 1 × 10^−14^).

**Figure 5.**
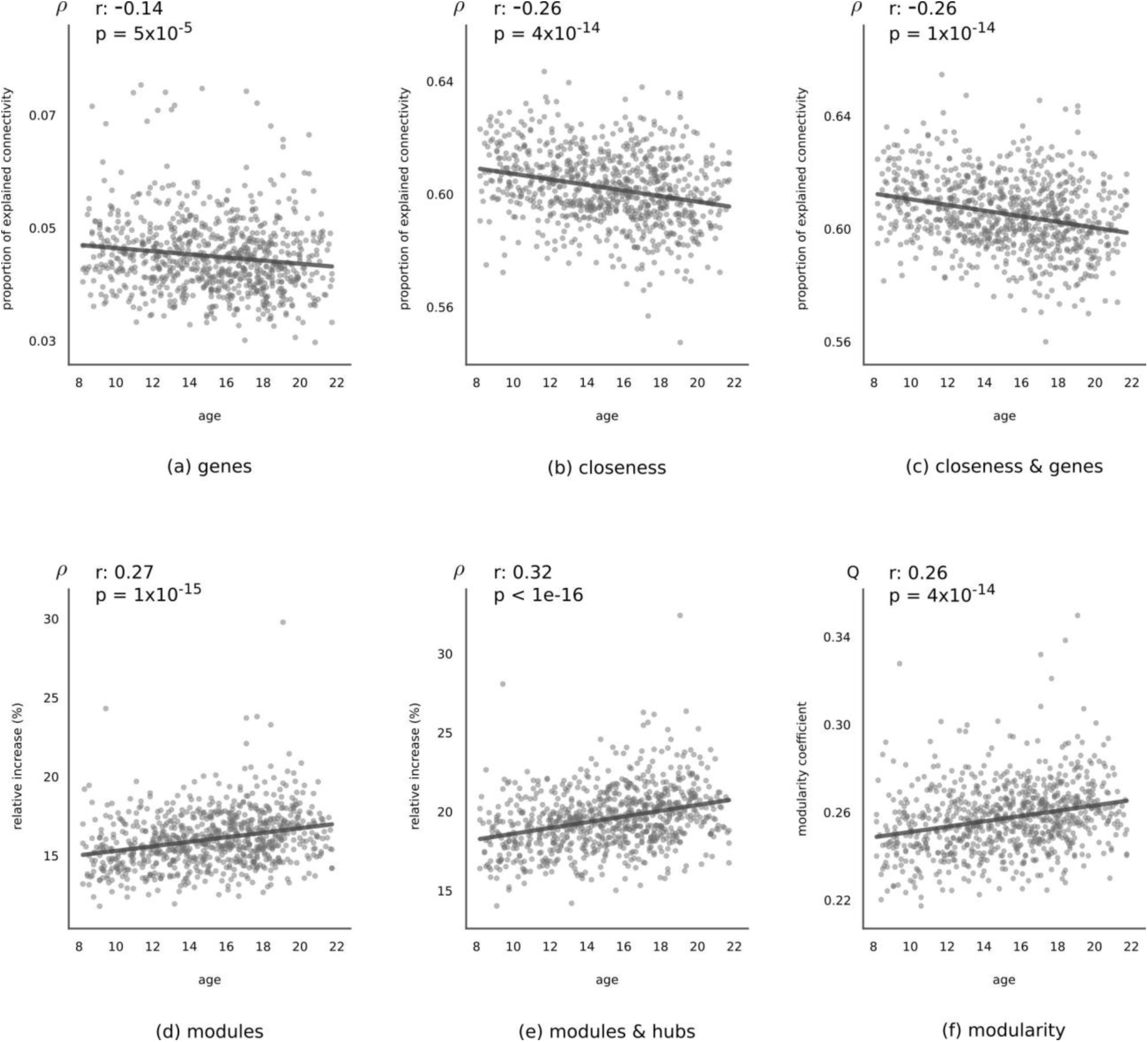
Developmental effects. (a,b,c) The proportion of explained connectivity as measured by ρ across ages, using (a) only genetic similarity factor, (b) only closeness factor, and (c) both. Pearson correlation coefficient (r) is given at the top of each figure. In all the three cases, explained connectivity shows a significant decrease over the course of development. The reverse is observed for the architectural specifications corresponding to the presence of (d) modules and (e) both modules and hubs. In (d) and (e), the relative increase in ρ compared to the base model (including both closeness and genetic similarity factors) is given. (f) Developmental change in the modularity coefficient (Q) that quantifies the extent that a network has a modular organization.

In contrast to the decreasing effect of the base model, the contribution of the architectural specifications in ρ was positively correlated with age (**Fig. 5d, e**). The relative change in ρ value as compared to the base model showed significant increase both for modular structure alone (Pearson correlation coefficient r = 0.27, p = 1 × 10^−15^) and for modular structure with hubs (Pearson correlation coefficient r = 0.32, p < 1 × 10^−16^). Additionally, the modularity coefficient (Q) that quantifies the extent that a network has a modular organization (that is, dense connections within modules, sparse connections between modules) [42] showed a significant increase by age (Pearson correlation coefficient r = 0.26, p < 4 × 10^−14^) (**Fig. 5f**).

We finally investigated how the relationship between structural network architecture and functional specialization of regions evolves during development. We calculated individual NMI values for each participant, by comparing alliance of regions determined by the structural modules and the predefined functional systems. We observed a significant positive correlation between individual NMI values and ages of participants (Pearson correlation coefficient r = 0.11, p = 0.002), indicating that the functional relevance of structural modules increases during development.

## DISCUSSION

We investigated how multiple factors contribute towards explaining observed streamline counts, and how their effects change over the course of development. We found a dominant contribution of brain geometry in explaining the observed streamlines, with regional gene expression profiles contributing less, albeit significantly. However, the brain network organization could not be fully accounted for by only considering geometric and genetic features of brain regions. Architectural specifications, which promote segregated modules and integrative hub regions, explained a significant proportion of the observed connectivity. Notably, the extent to which these architectural specifications contribute towards explaining structural connectivity increased during development, relative to the contribution of geometry and genes. Additionally, the functional relevance of the structural modules increased during development.

### Wiring Costs and Genes

The human brain develops under special sets of constraints reflected through its physical features. It is spatially embedded into a three-dimensional anatomical space and is subject to metabolic costs in forming axonal connections between regions [43], [44]. Many aspects of brain networks have been linked to such geometric and cost related constraints, via conservation principles such as minimization of wiring cost or energy consumption [45], [46]. Our results provide important insights into the contribution of geometry in explaining observed connectivity between regions, by quantifying the extent to which physical distance between regions explains the structural connectivity. The distance factor was the most dominant contributor in explaining the observed connectivity (**Fig. 3**). This finding is consistent with the major role of distance in the economy of human brain organization [15], [44], [47].

The effect of regional gene expression levels in the formation of axonal connections has recently begun to be explored in postmortem tissue samples [13]. Notably, in human brains, gene expression profiles define only subtle differences between regions of the neocortex [32], [36], with significant genetic dissimilarities among major sections, namely cortex, subcortex, and cerebellum. Our results were consistent with this fact, as the main genetic differences were observed among these three major sections (**Fig. 3**).

### Connectivity is More Than Wiring Costs and Genes

It is expected that the brain network organization cannot be fully accounted for by wiring costs alone [14], [15], [17], [18], [48]. By using our generative model, we quantified the gap between the observed streamline counts and what is expected from a base model considering only geometric and genetic features of the regions. The proportion of the connectivity explained by the base model (ρ=0.61) pointed to the presence of other factors accounting for the unexplained portion. One possible hypothesis is that brain structural connectivity is shaped in order to balance the trade-off between operating cost minimization and the adaptive value of the resulting organization [44].

### Distinctive Architecture of Brain Network

We demonstrated that the modular yet integrative organization of the brain, which has been commonly demonstrated for functional networks [4], is also evident in structural brain networks (**Fig. 4**). Our results supported the hypothesis that architectural specifications accompany geometric and genetic specifications in explaining the brain network organization [17]. Observed connectivity was explained best by a model that incorporates such architectural specifications related to the presence of modules and hubs (**Fig. 3**). Although our results demonstrated the importance of the architectural specifications, it is still unclear what other factors can be associated with these specifications. A promising future research direction can be the identification of molecular, as well as system-level features of the brain that correlate with this tendency to have a distinctive architecture.

The identified structural modules showed noticeable symmetry between hemispheres. Notably, the alliance of regions in forming structural modules highlights their functional correlates. One pair of symmetric modules (modules 1,4 in **Fig. 4**) was mainly defined by regions of frontal and cingulate cortex, encompassing the default mode network, cingulo-opercular system, and ventral attention system. One module that extended to both hemispheres (module 2) mainly consisted of regions of the visual system, including fusiform, inferior temporal, lateral occipital, and lingual. Homotopic visual areas are densely connected through callosal connections [49], possibly in order to facilitate the continuity of perception [50]. This may explain the presence of this bi-hemispheric module. The same module also included left and right cerebellar cortices, that are again densely connected to each other. The division of the temporal lobe between modules can be putatively associated with the separation between visual and auditory/language pathways. Superior and medial temporal cortices were assigned to modules 0 and 5 that included mainly parietal and subcortical regions. The connections between superior temporal cortex and inferior parietal cortex constitute an integral part of the speech processing system [51], [52]. On the other hand, the inferior temporal cortex was assigned to the bi-hemispheric visual module. The ventral stream of visual processing utilizes connections between visual cortex and inferior temporal cortex [53], [54].

Our analyses also demonstrated that the resulting modular organization of the structural brain network is significantly associated with emergent functional systems, supporting previous findings on the positive correlation between structural and fMRI-based functional connectivity [55], [56]. Elaborating the exact role of topological features, such as the presence of structural hubs, in facilitating links between structural modules and functional systems is a promising future research direction.

### Brain Network Becomes Even More Distinctive During Development

The human brain undergoes a protracted period of development from childhood to adulthood [57]–[59] that is assumed to be linked to the development of many sophisticated cognitive functions. Cognitive and behavioral changes occurring throughout childhood and adolescence [60] make these periods especially critical for investigations on changes of the white matter connectivity.

Developmental changes in the contribution of individual factors, as suggested by our results, presented an interesting picture of the developing brain during adolescence. The contribution of geometry in explaining streamline counts decreases over development, possibly implying that the links between wiring costs and the network organization of the brain weaken during development, with increasing emphasis on the adaptive value of the resulting organization. This possibility was further supported by the increasing effect of the architectural specifications that promote a modular organization (**Fig. 5**).

### Methodological Considerations

Several practical limitations of the current work should be considered when evaluating the presented results. Measuring structural connectivity using diffusion tensor imaging may suffer from the inability to accurately describe the regions with crossing fibers [61]. More advanced imaging techniques with higher resolutions [62] can be used in future studies where such data has been collected. Furthermore, diffusion based tractography has method-specific biases in fiber lengths, making the resulting connectivity matrices biased towards short-range connections [61], [63]. Therefore, the reported strong contribution of Euclidean distance in explaining the observed streamline counts may be partly attributed to the short-range bias of tractography algorithms. In order to demonstrate the robustness of the reported results to such confounding factors, we used consistency based thresholding [27] to prune possibly spurious connections and repeated our experiments with different network density (threshold) levels. The relative contribution of all factors remained the same across different density levels, suggesting high reliability of our results (**Supplementary Fig. S5**).

The geometric proximity of regions was computed using Euclidean distance. Other more sophisticated distance measures such as geodesic distance, defined within the white matter geometry, could also be used for the same purpose. However, this approach could lead to a circular argument when explaining connectivity by geometry, since white matter geometry needs to be defined by observed streamline connectivity.

The blurring effect caused by head motion during image acquisition may introduce a spurious increase in the contribution of geometry, which will possibly be higher in younger participants. In order to have results robust to motion effects, we have used the comprehensive quality assurance pipeline as described in [21]. The primary measurement of in-scanner head motion was mean relative volume-to-volume displacement as determined by rigid-body motion correction. According to this motion metric, our sample (mean: 0.46mm, std: 0.4mm) included images with “excellent” and “good” quality (please refer to Table 1 in Roalf et al. 2016). Nevertheless, more analyses can be done in future by including the amount of motion as a covariate, although which motion parameters to be included remains a current challenge.

In our analyses, we defined a generative model of structural connectivity using a parametric model (Dirichlet-Multinomial). Thus, the measure ρ that we used to calculate the proportion of the observed connectivity explained by the model, is a reliable measure for the same, as long as the selected parametric model is an appropriate choice for our dataset. In order to demonstrate that our model can reliably describe structural connectivity, we compared it with several nonparametric models, validating the plausibility of our parametric choices (**Supplementary Note S3**). We also showed that the reported results do not change when using a more conventional measure of performance. Instead of using the measure ρ, we repeated our experiments using root mean square error (RMSE) between empirical networks and the ones that are generated by our model. The contributions of factors were stable across different metrics, highlighting reliability of our results (see **Supplementary Note S9**).

Traditionally, hub regions are defined based on the degree, strength, centrality, or the participation coefficient distributions of the regions [64], [65]. For this reason, our definition of hubs based on certain architectural specifications may seem to be an unorthodox choice. We, therefore, showed that the identified hub regions have high participation coefficients and betweenness centrality (see **Supplementary Note S10**). This suggests that using certain architectural specifications to identify hub regions provides similar results as compared to traditional definitions.

The region-to-region genetic similarity was calculated from genome-wide expression levels of brain tissue samples from six different donors [33] of the Allen Brain Atlas [32]. The inter-regional genetic similarity factor was therefore same for all participants in our imaging sample. However, it is expected that the gene expression profiles, and hence the region-to-region similarities change over the course of development. The current expression profiles were computed using microarray data from adults (ages:24-57 years), which precludes a detailed analysis using our sample of young individuals (ages:8-22). This important limitation should be considered when interpreting our genetic findings. However, our work presents an important first step in studying the contribution of gene expression to brain organization, notwithstanding the fact that our data does not have *in vivo* characterization of the brain’s transcriptional distribution.

When studying the relationship between the *a priori* functional systems and the structural modules, we defined a fixed set of functional systems using results from a study [38] that had used a young adult sample (mean-age: 25.2 years, std: 2.5 years). In order to have more reliable interpretations regarding the relationship between functional systems and structural modules, however, future investigations should consider the fact that the functional organization of the brain also changes throughout development [2]. Defining functional systems for each participant individually, using a multimodal dataset including functional imaging modalities, is a possible future direction to take.

### A Generic Model for Future Neuroscience Studies

In this study, we considered effects of two intrinsic features, namely distance and gene expression, plus two architectural specifications corresponding to the presence of modules and hub regions. One advantage of the proposed methodology is the ease of inclusion of novel factors, including fiber lengths, cortical gyrification, and cytoarchitectonic or other histological properties of regions, providing a generic tool for future studies. Furthermore, probabilistic generative models can successfully represent different individual modalities of brain connectivity such as functional connectivity and can be used to model different modalities jointly. Multimodal connectivity is an especially exciting avenue of research to pursue.

Employing generative models of brain connectivity may further advance network neuroscience by providing a means of modeling joint and possibly nonlinear contributions of multiple factors in describing structural and functional characteristics of the brain network. This may enable development of new methodologies in effectively identifying alterations in connectivity patterns that have been shown to be present in clinical samples [66], between sexes [67], and during learning [68] or development [69].

## ACKNOWLEDGMENTS

This research was supported by the grant from the National Institutes of Health (R01-HD089390, PI: Ragini Verma), (R01-MH092862, PI: Ragini Verma and C-F Westin), (RO1-MH107235, PI: Ruben C. Gur).

## REFERENCES

[1] O. Sporns, D. R. Chialvo, M. Kaiser, and C. C. Hilgetag, “Organization, development and function of complex brain networks.,” Trends Cogn. Sci., vol. 8, no. 9, pp. 418–25, Sep. 2004.

[2] D. A. Fair, A. L. Cohen, J. D. Power, N. U. F. Dosenbach, J. A. Church, F. M. Miezin, B. L. Schlaggar, and S. E. Petersen, “Functional brain networks develop from a ‘local to distributed’ organization.,” PLoS Comput. Biol., vol. 5, no. 5, p. e1000381, 2009.

[3] D. Meunier, R. Lambiotte, and E. T. Bullmore, “Modular and hierarchically modular organization of brain networks.,” Front. Neurosci., vol. 4, p. 200, Jan. 2010.

[4] M. A. Bertolero, B. T. T. Yeo, and M. D’Esposito, “The modular and integrative functional architecture of the human brain.,” Proc. Natl. Acad. Sci. U. S. A., vol. 112, no. 49, pp. E6798–6807, Nov. 2015.

[5] P. Hagmann, L. Cammoun, X. Gigandet, R. Meuli, C. J. Honey, V. J. Wedeen, and O. Sporns, “Mapping the structural core of human cerebral cortex.,” PLoS Biol., vol. 6, no. 7, p. e159, Jul. 2008.

[6] E. Bullmore and O. Sporns, “Complex brain networks: graph theoretical analysis of structural and functional systems.,” Nat. Rev. Neurosci., vol. 10, no. 3, pp. 186–98, Mar. 2009.

[7] M. G. Mattar, M. W. Cole, S. L. Thompson-Schill, and D. S. Bassett, “A Functional Cartography of Cognitive Systems.,” PLoS Comput. Biol., vol. 11, no. 12, p. e1004533, 2015.

[8] G. Erus, H. Battapady, T. D. Satterthwaite, H. Hakonarson, R. E. Gur, C. Davatzikos, and R. C. Gur, “Imaging patterns of brain development and their relationship to cognition.,” Cereb. Cortex, vol. 25, no. 6, pp. 1676–1684, 2015.

[9] B. Tunç, B. Solmaz, D. Parker, T. D. Satterthwaite, M. A. Elliott, M. E. Calkins, K. Ruparel, R. E. Gur, R. C. Gur, and R. Verma, “Establishing a link between sex-related differences in the structural connectome and behaviour,” Philos. Trans. R. Soc. B Biol. Sci., vol. 371, no. 1688, 2016.

[10] A. E. Kahn, M. G. Mattar, J. M. Vettel, N. F. Wymbs, S. T. Grafton, and D. S. Bassett, “Structural Pathways Supporting Swift Acquisition of New Visuomotor Skills.,” Cereb. Cortex, vol. 27, no. 1, pp. 173–184, 2017.

[11] T. D. Satterthwaite, M. A. Elliott, K. Ruparel, J. Loughead, K. Prabhakaran, M. E. Calkins, R. Hopson, C. Jackson, J. Keefe, M. Riley, F. D. Mentch, P. Sleiman, R. Verma, C. Davatzikos, H. Hakonarson, R. C. Gur, and R. E. Gur, “Neuroimaging of the Philadelphia neurodevelopmental cohort.,” Neuroimage, vol. 86, pp. 544–53, Feb. 2014.

[12] A. Sotiras, J. B. Toledo, R. E. Gur, R. C. Gur, T. D. Satterthwaite, and C. Davatzikos, “Patterns of coordinated cortical remodeling during adolescence and their associations with functional specialization and evolutionary expansion.,” Proc. Natl. Acad. Sci. U. S. A., p. 201620928, Mar. 2017.

[13] P. Goel, A. Kuceyeski, E. LoCastro, and A. Raj, “Spatial patterns of genome-wide expression profiles reflect anatomic and fiber connectivity architecture of healthy human brain.,” Hum. Brain Mapp., vol. 35, no. 8, pp. 4204–18, Aug. 2014.

[14] J. A. Roberts, A. Perry, A. R. Lord, G. Roberts, P. B. Mitchell, R. E. Smith, F. Calamante, and M. Breakspear, “The contribution of geometry to the human connectome.,” Neuroimage, vol. 124, no. Pt A, pp. 379–93, Jan. 2016.

[15] D. Samu, A. K. Seth, and T. Nowotny, “Influence of wiring cost on the large-scale architecture of human cortical connectivity.,” PLoS Comput. Biol., vol. 10, no. 4, p. e1003557, Apr. 2014.

[16] J. A. Henderson and P. A. Robinson, “Relations between the geometry of cortical gyrification and white-matter network architecture.,” Brain Connect., vol. 4, no. 2, pp. 112–30, Mar. 2014.

[17] R. F. Betzel, A. Avena-Koenigsberger, J. Goñi, Y. He, M. A. de Reus, A. Griffa, P. E. Vértes, B. Mišic, J.-P. Thiran, P. Hagmann, M. van den Heuvel, X.-N. Zuo, E. T. Bullmore, and O. Sporns, “Generative models of the human connectome,” Neuroimage, vol. 124, no. Pt A, pp. 1054–64, Sep. 2015.

[18] M. Rubinov, “Constraints and spandrels of interareal connectomes.,” Nat. Commun., vol. 7, p. 13812, Dec. 2016.

[19] D. Heckerman, D. Geiger, and D. M. Chickering, “Learning Bayesian Networks: The Combination of Knowledge and Statistical Data,” Mach. Learn., vol. 20, no. 3, pp. 197–243, 1995.

[20] S. Mukherjee and T. P. Speed, “Network inference using informative priors.,” Proc. Natl. Acad. Sci. U. S. A., vol. 105, no. 38, pp. 14313–8, Sep. 2008.

[21] D. R. Roalf, M. Quarmley, M. A. Elliott, T. D. Satterthwaite, S. N. Vandekar, K. Ruparel, E. D. Gennatas, M. E. Calkins, T. M. Moore, R. Hopson, K. Prabhakaran, C. T. Jackson, R. Verma, H. Hakonarson, R. C. Gur, and R. E. Gur, “The impact of quality assurance assessment on diffusion tensor imaging outcomes in a large-scale population-based cohort,” Neuroimage, vol. 125, pp. 903–919, 2016.

[22] M. Jenkinson, C. F. Beckmann, T. E. J. Behrens, M. W. Woolrich, and S. M. Smith, “FSL,” Neuroimage, vol. 62, no. 2, pp. 782–90, 2012.

[23] S. Pieper, M. Halle, and R. Kikinis, “3D Slicer,” in IEEE International Symposium on Biomedical Imaging, 2004, pp. 632–635.

[24] C. Pierpaoli and P. J. Basser, “Toward a quantitative assessment of diffusion anisotropy.,” Magn. Reson. Med., vol. 36, no. 6, pp. 893–906, 1996.

[25] B. Fischl, M. I. Sereno, and A. M. Dale, “Cortical surface-based analysis. II: Inflation, flattening, and a surface-based coordinate system.,” Neuroimage, vol. 9, no. 2, pp. 195–207, Mar. 1999.

[26] R. S. Desikan, F. Segonne, B. Fischl, B. Quinn, B. Dickerson, D. Blacker, R. Buckner, A. Dale, R. Maguire, B. Hyman, M. Albert, and R. Killiany, “An automated labeling system for subdividing the human cerebral cortex on MRI scans into gyral based regions of interest,” Neuroimage, vol. 31, no. 2, 2006.

[27] J. Roberts, A. Perry, G. Roberts, P. Mitchell, and M. Breakspear, “Consistency-based thresholding of the human connectome,” Neuroimage, vol. 145, no. Pt A, pp. 118–129, 2017.

[28] S. W. Oh, J. A. Harris, L. Ng, B. Winslow, N. Cain, S. Mihalas, Q. Wang, C. Lau, L. Kuan, A. M. Henry, M. T. Mortrud, B. Ouellette, T. N. Nguyen, S. A. Sorensen, C. R. Slaughterbeck, W. Wakeman, Y. Li, D. Feng, A. Ho, E. Nicholas, K. E. Hirokawa, P. Bohn, K. M. Joines, H. Peng, M. J. Hawrylycz, J. W. Phillips, J. G. Hohmann, P. Wohnoutka, C. R. Gerfen, C. Koch, A. Bernard, C. Dang, A. R. Jones, and H. Zeng, “A mesoscale connectome of the mouse brain,” Nature, vol. 508, no. 7495, pp. 207–214, 2014.

[29] M. Hinne, T. Heskes, C. F. Beckmann, and M. A. J. van Gerven, “Bayesian inference of structural brain networks.,” Neuroimage, vol. 66, pp. 543–52, Feb. 2013.

[30] B. Tunç and R. Verma, “Unifying Inference of Meso-Scale Structures in Networks.,” PLoS One, vol. 10, no. 11, p. e0143133, Jan. 2015.

[31] H. Akaike, “A New Look at the Statistical Model Identification,” Trans. Autom. Control, vol. 19, no. 6, pp. 716–723, 1974.

[32] M. J. Hawrylycz, E. S. Lein, A. L. Guillozet-Bongaarts, E. H. Shen, L. Ng, J. A. Miller, L. N. van de Lagemaat, K. A. Smith, A. Ebbert, Z. L. Riley, C. Abajian, C. F. Beckmann, A. Bernard, D. Bertagnolli, A. F. Boe, P. M. Cartagena, M. M. Chakravarty, M. Chapin, J. Chong, R. A. Dalley, B. D. Daly, C. Dang, S. Datta, N. Dee, T. A. Dolbeare, V. Faber, D. Feng, D. R. Fowler, J. Goldy, B. W. Gregor, Z. Haradon, D. R. Haynor, J. G. Hohmann, S. Horvath, R. E. Howard, A. Jeromin, J. M. Jochim, M. Kinnunen, C. Lau, E. T. Lazarz, C. Lee, T. A. Lemon, L. Li, Y. Li, J. A. Morris, C. C. Overly, P. D. Parker, S. E. Parry, M. Reding, J. J. Royall, J. Schulkin, P. A. Sequeira, C. R. Slaughterbeck, S. C. Smith, A. J. Sodt, S. M. Sunkin, B. E. Swanson, M. P. Vawter, D. Williams, P. Wohnoutka, H. R. Zielke, D. H. Geschwind, P. R. Hof, S. M. Smith, C. Koch, S. G. N. Grant, and A. R. Jones, “An anatomically comprehensive atlas of the adult human brain transcriptome.,” Nature, vol. 489, no. 7416, pp. 391–9, Sep. 2012.

[33] AIBS, “Case qualification and donor profiles,” 2013. [Online]. Available: http://help.brain-map.org/download/attachments/2818165/CaseQual_and_DonorProfiles.pdf?version=1&modificationDate=1382051848013.

[34] AIBS, “Microarray data normalization,” 2013. [Online]. Available: http://help.brain-map.org/download/attachments/2818165/Normalization_WhitePaper.pdf?version=1&modificationDate=1361836502191.

[35] J. Richiardi, A. Altmann, A.-C. Milazzo, C. Chang, M. M. Chakravarty, T. Banaschewski, G. J. Barker, A. L. W. Bokde, U. Bromberg, C. Büchel, P. Conrod, M. Fauth-Bühler, H. Flor, V. Frouin, J. Gallinat, H. Garavan, P. Gowland, A. Heinz, H. Lemaître, K. F. Mann, J.-L. Martinot, F. Nees, T. Paus, Z. Pausova, M. Rietschel, T. W. Robbins, M. N. Smolka, R. Spanagel, A. Ströhle, G. Schumann, M. Hawrylycz, J.-B. Poline, and M. D. Greicius, “Correlated gene expression supports synchronous activity in brain networks.,” Science, vol. 348, no. 6240, pp. 1241–4, Jun. 2015.

[36] M. Hawrylycz, J. A. Miller, V. Menon, D. Feng, T. Dolbeare, A. L. Guillozet-Bongaarts, A. G. Jegga, B. J. Aronow, C.-K. Lee, A. Bernard, M. F. Glasser, D. L. Dierker, J. Menche, A. Szafer, F. Collman, P. Grange, K. A. Berman, S. Mihalas, Z. Yao, L. Stewart, A.-L. Barabási, J. Schulkin, J. Phillips, L. Ng, C. Dang, D. R. Haynor, A. Jones, D. C. Van Essen, C. Koch, and E. Lein, “Canonical genetic signatures of the adult human brain,” Nat. Neurosci., vol. 18, no. 12, pp. 1832–1844, 2015.

[37] M. Girvan and M. E. J. Newman, “Community structure in social and biological networks.,” Proc. Natl. Acad. Sci. U. S. A., vol. 99, no. 12, pp. 7821–6, Jun. 2002.

[38] J. D. Power, A. L. Cohen, S. M. Nelson, G. S. Wig, K. A. Barnes, J. A. Church, A. C. Vogel, T. O. Laumann, F. M. Miezin, B. L. Schlaggar, and S. E. Petersen, “Functional network organization of the human brain.,” Neuron, vol. 72, no. 4, pp. 665–78, 2011.

[39] S. Gu, F. Pasqualetti, M. Cieslak, Q. Telesford, A. Yu, A. Kahn, J. Medaglia, J. Vettel, M. Miller, S. Grafton, and D. Bassett, “Controllability of structural brain networks,” Nat. Commun., vol. 6, p. 8414, 2015.

[40] A. Alexander-Bloch, R. Lambiotte, B. Roberts, J. Giedd, N. Gogtay, and E. Bullmore, “The discovery of population differences in network community structure: new methods and applications to brain functional networks in schizophrenia.,” Neuroimage, vol. 59, no. 4, pp. 3889–900, Feb. 2012.

[41] V. D. Blondel, J.-L. Guillaume, R. Lambiotte, and E. Lefebvre, “Fast unfolding of communities in large networks,” J. Stat. Mech. Theory Exp., vol. 2008, no. 10, p. P10008, Oct. 2008.

[42] M. E. J. Newman, “Modularity and community structure in networks.,” Proc. Natl. Acad. Sci. U. S. A., vol. 103, no. 23, pp. 8577–82, Jun. 2006.

[43] S. Achard and E. Bullmore, “Efficiency and Cost of Economical Brain Functional Networks,” PLoS Comput. Biol., vol. 3, no. 2, p. e17, Feb. 2007.

[44] E. Bullmore and O. Sporns, “The economy of brain network organization.,” Nat. Rev. Neurosci., vol. 13, no. 5, pp. 336–49, May 2012.

[45] V. A. Klyachko and C. F. Stevens, “Connectivity optimization and the positioning of cortical areas.,” Proc. Natl. Acad. Sci. U. S. A., vol. 100, no. 13, pp. 7937–41, Jun. 2003.

[46] C. Cherniak, Z. Mokhtarzada, R. Rodriguez-Esteban, and K. Changizi, “Global optimization of cerebral cortex layout.,” Proc. Natl. Acad. Sci. U. S. A., vol. 101, no. 4, pp. 1081–6, Jan. 2004.

[47] D. B. Chklovskii, “Exact solution for the optimal neuronal layout problem.,” Neural Comput., vol. 16, no. 10, pp. 2067–78, Oct. 2004.

[48] M. Kaiser and C. C. Hilgetag, “Nonoptimal component placement, but short processing paths, due to long-distance projections in neural systems.,” PLoS Comput. Biol., vol. 2, no. 7, p. e95, Jul. 2006.

[49] S. Clarke and J. Miklossy, “Occipital cortex in man: organization of callosal connections, related myelo-and cytoarchitecture, and putative boundaries of functional visual areas.,” J. Comp. Neurol., vol. 298, no. 2, pp. 188–214, Aug. 1990.

[50] E. Genç, M. L. Schölvinck, J. Bergmann, W. Singer, and A. Kohler, “Functional Connectivity Patterns of Visual Cortex Reflect its Anatomical Organization.,” Cereb. Cortex, 2015.

[51] J. F. Démonet, F. Chollet, S. Ramsay, D. Cardebat, J. L. Nespoulous, R. Wise, A. Rascol, and R. Frackowiak, “The anatomy of phonological and semantic processing in normal subjects.,” Brain, vol. 115 (Pt 6, pp. 1753–68, Dec. 1992.

[52] G. Hickok and D. Poeppel, “The cortical organization of speech processing.,” Nat. Rev. Neurosci., vol. 8, no. 5, pp. 393–402, May 2007.

[53] A. Ishai, L. G. Ungerleider, A. Martin, J. L. Schouten, and J. V Haxby, “Distributed representation of objects in the human ventral visual pathway.,” Proc. Natl. Acad. Sci. U. S. A., vol. 96, no. 16, pp. 9379–84, Aug. 1999.

[54] P. Herath, S. Kinomura, and P. E. Roland, “Visual recognition: evidence for two distinctive mechanisms from a PET study.,” Hum. Brain Mapp., vol. 12, no. 2, pp. 110–9, Feb. 2001.

[55] C. J. Honey, R. Kötter, M. Breakspear, and O. Sporns, “Network structure of cerebral cortex shapes functional connectivity on multiple time scales.,” Proc. Natl. Acad. Sci. U. S. A., vol. 104, no. 24, pp. 10240–5, Jun. 2007.

[56] A. M. Hermundstad, D. S. Bassett, K. S. Brown, E. M. Aminoff, D. Clewett, S. Freeman, A. Frithsen, A. Johnson, C. M. Tipper, M. B. Miller, S. T. Grafton, and J. M. Carlson, “Structural foundations of resting-state and task-based functional connectivity in the human brain.,” Proc. Natl. Acad. Sci. U. S. A., vol. 110, no. 15, pp. 6169–74, Apr. 2013.

[57] E. L. Dennis and P. M. Thompson, “Mapping connectivity in the developing brain,” Int. J. Dev. Neurosci., vol. 31, no. 7, pp. 525–542, Nov. 2013.

[58] L. Q. Uddin, K. S. Supekar, S. Ryali, and V. Menon, “Dynamic Reconfiguration of Structural and Functional Connectivity Across Core Neurocognitive Brain Networks with Development,” J. Neurosci., vol. 31, no. 50, pp. 18578–18589, Dec. 2011.

[59] Z. M. Saygin, D. E. Osher, K. Koldewyn, R. E. Martin, A. Finn, R. Saxe, J. D. E. Gabrieli, and M. Sheridan, “Structural Connectivity of the Developing Human Amygdala,” PLoS One, vol. 10, no. 4, p. e0125170, Apr. 2015.

[60] A. W. Toga, P. M. Thompson, and E. R. Sowell, “Mapping brain maturation,” Trends Neurosci., vol. 29, no. 3, pp. 148–59, Mar. 2006.

[61] D. K. Jones, “Challenges and limitations of quantifying brain connectivity in vivo with diffusion MRI,” Imaging Med., vol. 2, no. 3, pp. 341–355, 2010.

[62] D. S. Tuch, R. M. Weisskoff, J. W. Belliveau, and V. J. Wedeen, “High Angular Resolution Diffusion Imaging of the Human Brain,” in Proceedings of the Annual Meeting of ISMRM, 1999.

[63] C. Reveley, A. K. Seth, C. Pierpaoli, A. C. Silva, D. Yu, R. C. Saunders, D. A. Leopold, and F. Q. Ye, “Superficial white matter fiber systems impede detection of long-range cortical connections in diffusion MR tractography.,” Proc. Natl. Acad. Sci. U. S. A., vol. 112, no. 21, pp. E2820–8, May 2015.

[64] O. Sporns, C. J. Honey, and R. Kötter, “Identification and classification of hubs in brain networks.,” PLoS One, vol. 2, no. 10, p. e1049, 2007.

[65] M. Rubinov and O. Sporns, “Complex network measures of brain connectivity: uses and interpretations.,” Neuroimage, vol. 52, no. 3, pp. 1059–69, Sep. 2010.

[66] M.-E. Lynall, D. S. Bassett, R. Kerwin, P. J. McKenna, M. Kitzbichler, U. Muller, and E. Bullmore, “Functional connectivity and brain networks in schizophrenia.,” J. Neurosci., vol. 30, no. 28, pp. 9477–87, Jul. 2010.

[67] M. Ingalhalikar, A. Smith, D. Parker, T. D. Satterthwaite, M. A. Elliott, K. Ruparel, H. Hakonarson, R. E. Gur, R. C. Gur, and R. Verma, “Sex differences in the structural connectome of the human brain.,” Proc. Natl. Acad. Sci. U. S. A., vol. 111, no. 2, pp. 823–8, Jan. 2014.

[68] D. S. Bassett, N. F. Wymbs, M. A. Porter, P. J. Mucha, J. M. Carlson, and S. T. Grafton, “Dynamic reconfiguration of human brain networks during learning.,” Proc. Natl. Acad. Sci. U. S. A., vol. 108, no. 18, pp. 7641–6, May 2011.

[69] K. Supekar, M. Musen, and V. Menon, “Development of large-scale functional brain networks in children.,” PLoS Biol., vol. 7, no. 7, p. e1000157, Jul. 2009.

